# Dihydrolipoamide dehydrogenase suppression induce human tau phosphorylation by increasing whole body glucose levels in a *C*. *elegans* model of Alzheimer’s Disease

**DOI:** 10.1101/220442

**Authors:** Waqar Ahmad

**Affiliations:** University of Queensland

## Abstract

The microtubule associated tau protein becomes hyperphosphorylated in Alzheimer’s disease (AD). While hyperphosphorylation promotes neurodegeneration, the cause and consequences of this abnormal modification are poorly understood. As impaired energy metabolism is an important hallmark of AD progression, we tested whether it could trigger phosphorylation of human tau protein in a transgenic *C. elegans* model of AD. We found that inhibition of a mitochondrial enzyme of energy metabolism, dihydrolipoamide dehydrogenase (DLD) resulted in elevated whole-body glucose levels as well as increased phosphorylation of tau. Hyperglycemia and tau phosphorylation were induced by either epigenetic suppression of the *dld-1* gene or by inhibition of the DLD enzyme by the inhibitor, 2-methoxyindole-2-carboxylic acid (MICA). Although the calcium ionophore A23187 could reduce tau phosphorylation induced by either chemical or genetic suppression of DLD, it was unable to reduce tau phosphorylation induced by hyperglycemia. While inhibition of the *dld-1* gene or treatment with MICA partially reversed the inhibition of acetylcholine neurotransmission by tau, neither treatment affected tau inhibited mobility. Conclusively, any abnormalities in energy metabolism were found to significantly affect the AD disease pathology.

## Introduction

Mitochondria plays an important role in cell bioenergetics and survival while mitochondrial dysfunction proposed as a key mediator of neurodegenerative diseases such as Alzheimer’s disease (AD) (1-3). In clinical studies, decline in cerebral glucose utilization, glucose-dependent energy production and associated enzyme’s activities such as pyruvate dehydrogenase (PDH) and α-ketoglutare dehydrogenase (KGDH) are observed in AD patients, which highlights the importance of glucose energy metabolism in disease progression (4-8).

Abnormal glucose levels could be linked to formation of neurofibrillary tangles (NFTs), aggregates of hyper-phosphorylated tau protein that is diagnostic features of advanced AD (9, 10). Tau regulates, maintains and stabilizes microtubule assembly under normal physiological conditions. Under such conditions, an average of 2-3 phosphate groups is present on each tau molecule, whereas in the AD brain the level of tau phosphorylation is 3-4 times higher (11, 12). Hyperphosphorylation leads to self-aggregation of tau filaments and activation of caspase-3 *in vitro*. *In vivo,* hyperphosphorylation is associated with tissue deterioration and cell death (13, 14).

Tau phosphorylation is negatively regulated by another post-translational modification *O*-GlcNAcylation of serine or threonine residues by UDP-GlcNAc at sites that overlap phosphorylation sites (15, 16). Given that UDP-GlcNAc biosynthesis is impaired by glucose catabolism, glucose catabolism could also cause reduced *O*-GlcNAcylation. As a result, down regulation of O-GlcNAc by low glucose levels is expected to promote tau hyperphosphorylation (17, 18). Although it has not yet been determined whether low glucose levels are a cause or consequence of AD, the relationship between glucose levels, *O-*GlcNAcylation and tau hyperphosphorylation provides a plausible causal mechanism. Indeed, a drop in glucose energy metabolism has been proposed to minimize AD-associated pathology, but the authors of the study proposed a mechanism in which suppression of energy metabolism compensates for the reduced nutrient and oxygen supply in the neuronal microenvironment (19). Moreover, improved prognosis against AD symptoms observed in several model systems under caloric restriction or reducing glucose-dependent energy production also supports the hypothesis that decrease in glucose metabolism may be a protective measure against AD (20-22).

Dihydrolipoamide dehydrogenase (*dld*) encodes a core metabolic enzyme and a part of two major enzyme complexes that contribute directly to glucose energy metabolism, PDH and KGDH (23). Remarkably, four single nucleotide variants of the *dld* gene are associated with increased risk of late-onset AD, though how each variant specifically affects DLD activity is still unknown (24). Interestingly, in brains from Swedish AD patients with the APP670/671 mutation *Aβ* was overproduced and all subunits of KGDH were suppressed except DLD, indicating that different subunits respond differentially to AD (25). In simple organisms *dld* gene suppression improves stress resistance in insects and increases lifespan in *C. elegans* (26, 27), suggesting that a therapeutic intervention that either suppressed *dld* gene expression or DLD enzyme activity could inhibit the symptoms of AD.

5-methoxyindole-2-carboxylic acid (MICA) is a chemical inhibitor of DLD enzyme activity whose effects can be reversed by calcium ionophore-A23187 (CaI), which increases intracellular calcium levels (28, 29). In the present study, we diminished the activity of DLD by RNAi or by exposure to MICA in transgenic *C. elegans* expressing human tau. This allowed us to investigate the possible role of glucose energy metabolism in relation to DLD actvity on tau phosphorylation. We find that increase in whole-body glucose levels that consequently, increases phosphorylation of several serine and threonine residues of tau that are important in NFTs formation after *dld* inhibition could be revrsed by CaI.

## Materials and methods

### Nematode strains

The wild-type strain was N2 (Bristol), whereas transgenic strain VH255 (hdEx82[F25B3::tau352 (WT) + pha-1 (+)] expressing human fetal 352aa CNS tau was used to assess the phosphorylation on tau and maintained as described earlier (30). Although our tau transgenic strain express fetal tau, the sites and magnitude of phosphorylation of fetal tau is similar to PHF-tau in AD (31).

### Culture conditions

Mixed-stage cultures of *C. elegans* were maintained on nematode growth medium (NGM) seeded with *E. coli* OP50 at 20°C. Assays utilised synchronised cultures, which were obtained by harvesting eggs from gravid hermaphrodites by exposing them to a freshly prepared alkaline bleach solution (0.75N NaOH + 1.5N NaOCl). Exposure was for five minutes at room temperature followed by rinsing 3 times with M9 buffer (6 g/L Na_2_HP0_4_; 3 g/L KH_2_P0_4_; 5 g/L NaCl; 0.25g/L MgSO_4_ •7H_2_0) and centrifugation at 1100 rpm for 1 minute at room temperature. Supernatant was discarded, and egg’s pellets were placed freshly made NGM plates at 25°C overnight for hatching. Next day, the resulting L1 larvae were shifted to their respective plates. Worms were monitored by visual observation under a microscope, and/ or quantified using the WormScan (32).

### dld-1 suppression by RNAi

The *E. coli* strain SJJ_LLC1.3 (Source Bioscience) was used to suppress the *dld-1* gene in each strain (33). Briefly, feeding bacterial strain was cultured in LB media containing 100μg/ mL ampicillin overnight in shaker at 37°C and kept at 4°C for further usage. 300 μL of feeding bacteria was transferred to NGM plates containing 100μg/ mL ampicillin and 1mM IPTG. Plates were kept at 25°C overnight to grow bacterial cultures. Synchronised worms were shifted to RNAi feeding plates and kept at 20°C for 48 hours before using for any assay.

### Chemical inhibition of dld complex in C. elegans using MICA

As MICA is partially soluble in water, stock solution of MICA was prepared by adding drop wise 5N sodium hydroxide till MICA becomes clearly soluble in water.

### Acetylcholine induction assay

Worms with and without *dld-1* inhibition were incubated in the presence of 1mM aldicarb an acetylcholinesterase inhibitor (34) and scored for aldicarb-induced paralysis. Active worms were counted every 30 minutes till last worm became paralyzed.

### Thrashing assay

Thrashing assays were counted in M9 buffer under microscope. Briefly, at least 10 worms were randomly picked from each group and their thrashing rates were counted in a time interval for 10 seconds at 20°C.

### Western Blotting

To detect phosphorylation on tau protein, synchronised worms were incubated at 20°C. L4 worms (day 3) were collected and washed three-four times to completely remove the bacterial traces. Half of the washed worms were added in 400μl cold RIPA buffer containing proteinases and phosphatases. While remaining half of the worms were shifted to new plates containing 75μM FUDR to restrict progeny production and collected on 7^th^ day. Worms were subjected to sonication and lysate was collected for further protein analysis. Total protein in the supernatant was measured using Pierce Coomassie (Bradford) protein assay kit (Thermo Scientific) on Nano drop. From each sample, 80-100 μg of total protein was precipitated with acetone and dissolved in Novex^®^ Tricine SDS sample buffer (LC1676, Invitrogen) by heating to 99°C for 5 minutes. 50-80ng of whole protein was dissolved in SDS sample buffer and subjected to standard 12.5% gel electrophoresis. Proteins were transferred to nitrocellulose membrane for 60 min at 17V for semi-dry transfer. Membranes were blocked in 5%BSA in 0.1% TBST containing 0.1% Tween 20. A number of antibodies were used to assess total and site specific phosphorylated tau. The antibodies used for tau were, Tau-5 (ab80579, abcam), HT7 (MN1000, Thermo Scientific), phosphor S198 (ab79540, abcam), phosphor S235 (NB100-82241, Novus Biologicals), phosphor S262 (79-152, Prosci), phosphor S396 (710298, Novex Lifetecnologies), AT8 (S202, MN1020, Thermo Scientific), and AT180 (T231, MN1040, Thermo Scientific). Anti-actin antibody (2Q1055, abcam) was used as reference control. Anti-mouse IgG alkaline phosphatase antibody produced in goat (A3562, Sigma), and anti-rabbit IgG alkaline phosphatase antibody produced in goat (A3687, Sigma were used as secondary antibody at 1:10000 dilution in 1% PBS in 0.1% TBST. 2ndary antibody staining was done for 1 hour at room temperature. After washing the membrane with TBST, the proteins were detected using BCIP/ NBT substrate system (Sigma) or BCIP/ NBT kit (002209) from Lifetechnologies dissolved in 1M Tris (ph 9.0).

## Determination of total glucose

Total glucose concentration in whole worm lysate was determined using Sigma glucose assay kit (GAGO20) under described method. Briefly, transgenic tau VH255 synchronised worms were placed on NGM plates containing different MICA and/ or glucose concentrations at 20°C. Day 3 worms were collected and washed with M9 buffer to remove any bacterial contents. Worm pellets were dissolved in ice cold RIPA buffer, flash frozen in liquid nitrogen, and placed at −80° C for further analysis. Worms were sonicated and lysate was collected after centrifugation at 10000rpm at 4°C. Pellet was discarded. Total protein content was measured using Pierce Coomassie (Bradford) protein assay kit (Thermo Scientific) on Nano drop, and used as normalisation factor. Same lysate was used for total glucose determination in mg and converted to mM (1mg/ dl glucose = ~5.5mM).

### Reversal of MICA-mediated effects

Transgenic worm VH255 were synchronized and placed at NGM plates containing *dld-1* RNAi and/ or 5mM MICA for 24 hours at 20°C. L2 staged worms were then shifted to NGM plates containing 35μM calcium ionophore A23187 (CaI) with and without MICA/ *dld-1* RNAi. Whole body proteins were extracted from L4 worms and proceed further for analysis.

### Statistical analysis

Mean, standard deviation and *p* values were calculated using GraphPad (San Diego, CA) prism 6.0.3. Differences between treated and untreated worm strains were analysed for statistical significance by independent student’s t-test or one-way Anova. A *p* value less than 0. 05 was considered statistically significant.

## Results

In this study we used the *C. elegans* strain that express human fetal tau in neurons. As we are interested to find association between impaired energy metabolism and AD, here in this study we investigated the effects of suppression of core metabolism enzyme DLD on tau-mediated toxicity in *C*. elegans. Tau become phosphorylated in AD that is a complex phenomenon. While hyperphosphotylation is generally associated with toxicity of tau, specific sites may be differentially phosphorylated and may contribute disproportionately to toxicity. Previous studies revealed that Ser-46, Ser-198, Ser-199, Ser-202, Thr-205, thr-212, Thr-231/Ser-235, Ser-262/Ser-356, Ser-396, Ser-404, and Ser-422 are among critical sites in AD and hyperphosphorylation on these residues hinders its binding with microtubules and promote NFTs formation(9, 12, 35-37). In this study we investigated the effect of *dld-1* suppression on tau toxicity and phosphorylation on several critical sites/ epitopes to draw any association between glucose energy metabolism and tau phosphorylation in AD.

To check whether inhibition of either the *dld-1* gene or DLD protein affects phosphorylation of tau, phosphorylation on specific tau epitopes was analysed using site-specific phospho antibodies on day 3 and day 7 after *dld-1* suppression/ inhibition. The HT7 antibody that represent total tau (independent of phosphorylation) showed that inhibition of *dld-1* does not changed overall tau levels in worms until day 7. We observed a significant increase in phosphorylation pSer-198 (27.1 fold), pSer-235 (12.85 fold), and pSer-262 (12.3 fold) in MICA (DLD inhibited) treated samples on day 3 that become more prominent on day 7 however, at this time transgenic control also showed phosphorylation. Although on day 3, worms fed with *dld-1* RNAi did not showed any significant increase in phosphorylation on any of the sites relative to the untreated transgenic control, by day 7 there was a significant increase in phosphorylation on Ser-198 (4.01 fold), Ser-235 (3.43 fold) and Ser-262 (7.71 fold) epitopes. The most prominent difference in the response to MICA relative to *dld-1* RNAi is a difference in the speed of phosphorylation. This is most likely due to direct inhibition of the enzyme being faster than a decrease in gene expression that requires protein turnover to occur before the effect can be observed.

DLD is a part of important enzyme complex PDH that links between glycolysis and Kreb’s cycle. Previous studies showed that inhibition of DLD through MICA increased intracellular glucose levels (28, 38). Our aim was to evaluate whether this increase in tau phosphorylation in MICA treated worms is linked with increased glucose levels. To assess this, we fed worms with different concentrations of MICA (ranging from 5mM – 0.625 mM) and glucose (4% −0. 25%) separately in NGM. We found that both MICA and glucose increased phosphorylation of tau dose dependantly. Concentrations of MICA 0.625, 1.25, 2.5 and 5mM induce phosphorylation on pSer-198 by 5.1, 6.6, 9.4 and 16.8 fold, and on pSer-235 by 1.5, 1.7, 2.1, and 4.9 fold, respectively (Fig 2A). Meanwhile glucose concentrations of 0.25, 0.5, 1, and 4% induce phosphorylation on pSer-198 by 7.4, 10.1, 13.6 and 15.6 fold, and on pSer-235 by 1.5, 1.8, 1.8, and 3.1 fold, respectively (Fig 2B).

**Figure 2:**
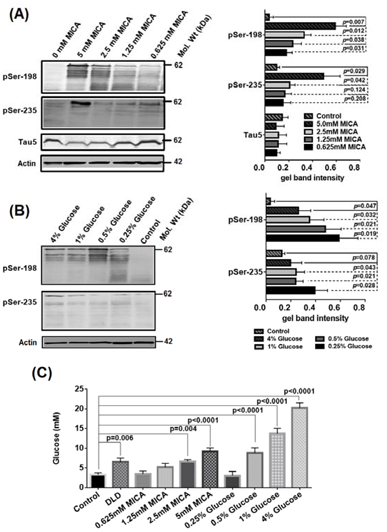
MICA induce tau phosphorylation dose dependantly that can be attributed to induced intracellular glucose levels. (A) Tau expressing worms were synchronised and shifted to 0, 5, 2.5, 1.25, and 0.625mM concentrations of MICA. B) To check whether glucose modulate tau phosphorylation, worms were fed with different concentrations of glucose (0.25, 0.5, 1, and 4%). Proteins were extracted after 3 day incubation and tau phosphorylation was detected. Tau phospho antibodies specific to Ser-198 and Ser-235 were used to detect any changes in tau phosphorylation. Anti-actin antibody was used as refence control. Quantification of gel bands from each row was done using GelQuantNET. Graph are shown from two independent trials. C) Determination of whole body glucose levels in tau transgenic worms. Synchronised worms were shifted to different concentrations of glucose and MICA or fed with *dld-1* RNAi. Glucose concentrations were mesured and normalized against extracted protein after 3 days incubation. Glucose concentrations were measured in mg and then converted to mM (1mg/ dl glucose = ~5.5mM). Graph are shown from three independent trials. Bars = Mean±SD.

Glucose quantification in worms fed with *dld-1* RNAi, MICA and glucose showed significant increase in whole body glucose levels (Fig 2C). The glucose concentration in untreated VH255 control worms was 3.35±0.95 mM, while addition of 5mM MICA in NGM resulted in elevated intracellular glucose concentration 9.12±0.91 mM (p<0.0001). Same as MICA, addition of glucose also increased intracellular glucose cencentration where 0.5% glucose induced intracellular glucose as of 5mM MICA (8.85±1.26, p=0.729). *Dld-1* suppression by RNAi also resulted in increase in intracellular glucose that was significantly higher than control (6.47±1.05, p=0.008) and almost equal to 1.25Mm MICA treatment (5.23±0.93, p=0.198).

Inhibition of DLD was found to induce phosphorylation on all tau epitopes in this study (Fig 1 and 2). As inhibition of DLD activity can be reversed using CaI (29), we assessed whether CaI was able to reverse this induction in phosphorylation in *dld-1* suppressed/ DLD inhibited worms. Treatment with CaI reduced tau phosphorylation on pSer-198, pSer-262, and AT180 epitope by 2.9, 3.1, and 5.8 fold in worms fed with *dld-1* RNAi, and 3.4, 2.6, and 11.6 fold in MICA fed worms, respectively. CaI as it does not show any difference in tau phosphorylation profile when compared to control.

**Figure 1:**
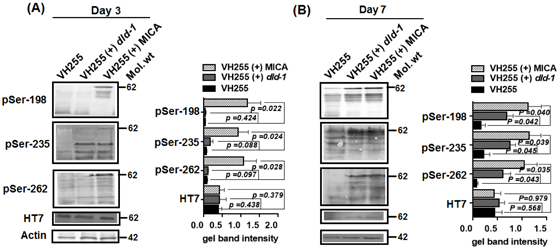
Effect of dld inhibition on tau phosphorylation. Worms were either fed with *dld-1* RNAi or with 5mM MICA after synchronization. Proteins were extracted after three days or seventh day of feeding to check long term effect of *dld-1* suppression. Extracted proteins were run on 10% SDS PAGE and tau was detected by using non-phosphorylated antibody HT-7, and site specific phosphor antibodies Ser-198, Ser-235 and Ser-262. Anti-actin antibody was used as reference control. A) Antibody staining of worm protein extracted after three days. B) Antibody staining of worm protein extracted after seventh day. Quantification of gel bands from each row was done using GelQuantNET. Graphs are shown from two independent trials. Bars = Mean ±SD.

In our study we observed that exposure to a diet supplemented with glucose resulted in induction of tau phosphorylation (Fig 1 and 2) whereas treatment with CaI reduced tau phosphorylation in presence of DLD inhibition (Fig 3). Our aim was to evaluate whether treatment with CaI reduced the phosphorylation of tau in the presence of high glucose. We found, however, that CaI was not able to minimize phosphorylation in worms fed with either 1% or 4% glucose (Fig 4).

**Figure 3:**
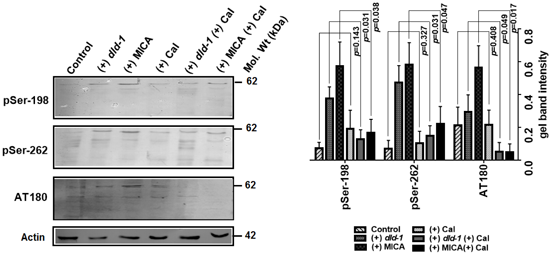
Effect of CaI on tau phosphorylation in presence of *dld-1* suppression and DLD inhibition. Tau phosphorylation induced by disruption of DLD may be reduced by CaI. Synchronised tau expressing worms were shifted for 24 hours to 35μM CaI plates with or without *dld-1* gene inhibition by RNAi or DLD enzyme inhibition by 5mM MICA. Whole body protein was extracted from worms after 7 days and evaluated for phosphorylation on respective tau epitopes using antibodies anti Ser-198, Ser-262, AT180. Quantification of gel bands from each row was done using GelQuantNET. Graph are shown from two independent trials. Bars= Mean±SD

**Figure 4:**
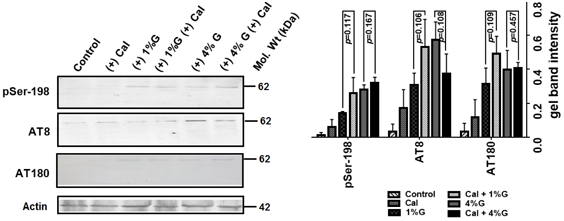
Induction of tau phosphorylation by glucose could not be countered by CaI. Tau expressing worms were fed with 1% or 4% glucose in the presence or absence of 35μM CaI. Whole body protein was extracted after 3 days and subjected to immunoblotting using tau phospho epitopes namely Ser-198, AT8 and AT180. Quantification of the intensities of each band relative to actin in each lane was carried out using GelQuantNET software. Graph are shown from two independent trials. Bars= Mean±SD

In this study we revealed that *dld-1* suppression and DLD inhibition increased tau phosphorylation that could result in induced tau pathology. Transgenic tau expression in worms was found negatively affects the synaptic transmission of cholinergic neurons and induced resistance against acetylcholine esterase (AChE) inhibitor aldicarb when compared to wild type (39). To check whether *dld-1* suppression or inhibition could affect acetylcholine neurotransmission, we treated the worms with aldicarb. In our study, when treated with aldicarb, wild type N2 worms become paralysed within 120 minutes when compared to tau expressing VH255 strain that took 180 minutes to become paralysed (p<0.0001). We found earlier paralysis in tau expressing worms compared to transgenic control in the presence of aldicarb when fed with either *dld-1* RNAi (150 min, p= 0.0131) or MICA (150 min, p=0.0272). The reduction in paralysis time by *dld-1* RNAi was not significantly different from suppression of enzyme activity by MICA (p=0.557). Although suppression of *dld-1* was able to reduce aldicarb resistance in transgenic worms, it was not up to the wild type (*dld-1* RNAi, 120 vs 150 min, p=0.0431; MICA, 120 vs 150 min, p=0.0038). Thus, restoration of normal acetylcholine signalling is the only AD-like symptom that can be alleviated by *dld-1* suppression in the *C. elegans* tau model of AD.

Improvements in acetylcholine neurotransmission could affect the worm locomotion. When submerged in liquid, *C. elegans* will move vigorously, but will be unable to gain traction as they would on a solid surface. This is referred to as thrashing. Expression of human tau in *C. elegans* resulted in reduced thrashing rates (30). Suppression of either the *dld-1* gene by RNAi or the DLD enzyme by exposure to MICA failed to restore normal movement. We monitored the thrashing rates till day seven and observed no improvement. Thrashing rates decreased gradually over the course of the experiment irrespective of treatment or genotype.

## Discussion

In this study we examined the effect of suppression of *dld-1* gene expression or chemical inhibition of the DLD enzyme activity on phosphorylation of the human tau protein in a *C. elegans* model of Alzheimer’s disease. As inhibition of either the dld gene or enzyme activity significantly disrupts glucose catabolism, we also explored the relationship between glucose levels and tau phosphorylation. We then used movement and neural function assays to monitor the relationship between tau phosphorylation and pathology. DLD is a subunit of PDH, an enzyme complex that regulates glucose catabolism at the interface between glycolysis and the TCA cycle (40). MICA is a chemical inhibitor of DLD that we used in this study. In *H. volcanii* and in mouse models, MICA does not exclusively inhibit PDH as described previously (28, 41), but also dehydrogenases of other substrates including succinate, lactate and alpha-ketoglutarate. Therefore, to complement our studies with MICA, we also carried out specific epigenetic suppression of the *dld-1* gene.

Inhibition of DLD and subsequently other mitochondrial dehydrogenase may lead to rise in glucose concentrations by reducing the carbon of pyruvate into the citric acid cycle (42) and might causing hyperglycaemia like stage in worms. Hyperglycaemia increases phosphorylation of tau in mouse models of diabetes (43-45). As DLD is a subunit of PDH, an enzyme that regulates glucose catabolism, we reasoned that inhibition of either DLD enzyme activity or expression of the *dld-1* gene would decrease the glycolytic catabolism of glucose (42) causing a rise in glucose concentration in *C. elegans.* In our study, inhibition of DLD by MICA resulted in an increase in phosphorylation at multiple sites of the tau protein. The response to *dld-1* gene suppression was slower, but after 7 days, a similar increase in phosphorylation at multiple sites of the tau protein occurred (Fig 1). The response to MICA was dose-dependent and as exposure to MICA and *dld-1* gene suppression both caused whole-body levels of glucose to increase, the effect of glucose on phosphorylation of tau was also determined (Fig 2). Like the situation in response to MICA exposure, the response to glucose was also seemed dose dependently.

CaI is reported to act as a MICA antagonist (46, 47). CaI is known to decrease the phosphorylation of the PDH enzyme complex and regulate glucose transport to cells (48-52). In our study, CaI alone did not affect the overall pattern of tau phosphorylation but it did prevent induction of phosphorylation by either *dld-1* RNAi or exposure to MICA (Fig 3). Interestingly, CaI failed to reduce tau phosphorylation in the presence of high glucose (Fig 4).

The failure to prevent the phosphorylation due to elevated glucose in the diet was surprising given the ability of CaI to prevent phosphorylation due to inhibition of the *dld-1* gene or DLD enzyme. This result indicates either that the phosphorylation occurs by different mechanisms or that the effect of CaI is mediated downstream of DLD, but upstream of glucose in a response pathway.

Expression of tau in *C. elegans* results in impaired cholinergic neurotransmission (39, 53) and progressive aged-dependent uncoordinated movement. Resistance against acetylcholinesterase inhibitor “aldicarb”, may be due to either a pre- or post-synaptic defect (54). Treatment with aldicarb causes more severe muscle contraction and early paralysis in wild type *C. elegans* compared to tau transgenic strains. Although the mechanism is unknown, both pre and post-synaptic defects observed in tau transgenic control were partially reversed by *dld-1* inhibition either using RNAi or MICA (Fig 5). Improvement in acetylcholine neurotransmission could improve worm’s uncoordinated movement. We found that *dld-1* inhibition did not reduce uncoordinated movement and that the thrashing rates decreased gradually with age (Fig 6). These results are consistent with a previously published report that attributed impaired movement to microtubules being “overstabilzed” by the expression of transgenic tau (30). As both *dld-1* gene suppression and treatment with MICA increase phosphorylation of tau without affecting the amount of tau protein, the ameliorative effect seems to be independent of tau or the general phosphorylation state of the protein. These results are in agreement with previous findings in which early staged-synaptic regulation and uncoordinated movement was independent of tau phosphorylation (30, 55).

**Figure 5:**
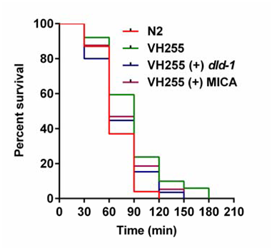
Aldicarb sensitivity assay in wild type and tau expressing worms. Time-dependent paralysis of worms on 1mM aldicarb fed with and without *dld-1* RNAi or 5mM MICA. Inhibition of either *dld-1* gene expression or DLD enzyme activity by MICA modulates acetylcholine neurotransmission. Paralysis curves were compared using Log-rank method. Results were derived from three independent experiments.

**Figure 6:**
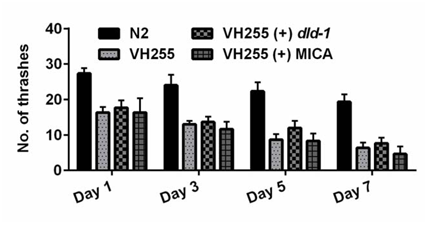
Inhibition of either *dld-1* gene expression or DLD enzyme activity does not restore thrashing rates of tau expressing worms. Quantification of thrashing rates in tau expressing worms (VH255) with and without *dld-1* gene or DLD enzyme suppression. Thrashing rates were quantified every second day from the start of the experiment until day 7 and compared with wild type (N2) worms that did not express tau.

In conclusions, we found that impairment of the enzyme DLD, which is involved in glucose energy metabolism, significantly induces phosphorylation of the tau protein. The outcome was the same whether impairment was by post-transcriptional silencing of the *dld-1* gene or by inhibition of the DLD enzyme by MICA. The phosphorylation induced by either method was reversible by the CaI. Both MICA and *dld-1* gene suppression resulted in elevated levels of whole-body glucose. While elevated glucose also resulted in increased phosphorylation of the tau protein, this effect could not be reversed by exposure to the CaI. Our results suggest that glucose-mediated phosphorylation occurs by a different mechanism or that it is mediated downstream of Ca^2+^ signalling.

